# Nested biofabrication: Matryoshka-inspired Intra-embedded Bioprinting

**DOI:** 10.1101/2023.09.28.560028

**Authors:** Mecit Altan Alioglu, Yasar Ozer Yilmaz, Yogendra Pratap Singh, Momoka Nagamine, Nazmiye Celik, Myoung Hwan Kim, Vaibhav Pal, Deepak Gupta, Ibrahim T. Ozbolat

## Abstract

Engineering functional tissues and organs remains a fundamental pursuit in biofabrication. However, the accurate constitution of complex shapes and internal anatomical features of specific organs, including their intricate blood vessels and nerves, remains a significant challenge. Inspired by the Matryoshka doll, we here introduce a new method called ‘Intra-Embedded Bioprinting (IEB),’ building upon existing embedded bioprinting methods. We used a xanthan gum-based material, which served a dual role as both a bioprintable ink and a support bath, due to its unique shear-thinning and self-healing properties. We demonstrated IEB’s capabilities in organ modelling, creating a miniaturized replica of a pancreas using a photocrosslinkable silicone composite. Further, a head phantom and a Matryoshka doll were 3D printed, exemplifying IEB’s capability to manufacture intricate, nested structures. Towards the use case of IEB and employing innovative coupling strategy between extrusion-based and aspiration-assisted bioprinting, we developed a breast tumor model that included a central channel mimicking a blood vessel, with tumor spheroids bioprinted in proximity. Validation using a clinically-available chemotherapeutic drug illustrated its efficacy in reducing the tumor volume via perfusion over time. This method opens a new way of bioprinting enabling the creation of complex-shaped organs with internal anatomical features.

## 1. Introduction

In recent years, the merging of additive manufacturing and bioengineering has initiated the innovative field of three-dimensional (3D) bioprinting. This technology has the potential to revolutionize tissue engineering, regenerative medicine, and personalized healthcare by enabling the deposition of bioinks layer by layer to form 3D living constructs ^[1]^. However, a significant hurdle in the path of successful bioprinting involves the deformation of the extruded soft bioinks due to the influence of gravity, resulting in compromised print fidelity. The absence of a solid support complicates the layer-by-layer bioprinting of most soft bioinks, which may not solidify rapidly or with adequate rigidity to maintain their structural integrity throughout the bioprinting process. Researchers have addressed this issue by developing innovative strategies like photo-crosslinkable, temperature-sensitive, and rheologically-modified bioinks, aimed at mitigating the impact of gravity and facilitating 3D bioprinting in air without a support ^[2–4]^. Nevertheless, these approaches often necessitate trade-offs in terms of the depleted inherent biological properties of the bioink, aiming to strike a balance with the material properties necessary for optimal printability.

Recently, embedded bioprinting has emerged as a promising approach to tackle this challenge, where complex tissue structures are fabricated directly within a reservoir containing a supportive matrix, efficiently making print fidelity independent from the bioink’s gelation time and crosslinking methods. Benefitting from the pre-existing matrix as a support, bioinks with low viscosity and mechanical strength can be stably bioprinted into predefined patterns ^[5]^. This method can be envisioned as a construction chamber filled with a supporting material. Inside this chamber, cells, spheroids, cell-laden hydrogels, and other substances can be precisely deposited on demand ^[6,7]^. A fundamental engineering challenge in embedded bioprinting is developing a support bath material capable of simultaneously maintaining the shape of the bioprinted structures during extrusion and curing, while also facilitating the extruder needle’s movement through the support bath during bioprinting. Consequently, the support bath must have yield-stress behavior, akin to a Bingham plastic or Herschel–Bulkley fluid ^[6]^. In this context, the support material functions as solid until adequate shear stress (known as yield stress) is applied. In embedded bioprinting, the movement of the needle inside the support bath produces a shear stress around the needle, which is greater than the yield stress of the bath material. At this juncture, the material transitions from a solid to a fluid-like state, allowing the smooth motion of the needle as well as the successful deposition of the bioink. As soon as the needle moves away, the bath material immediately turns back to its solid form in the absence of the shear stress, thereby locking the extruded bioink in its position until it solidifies, counteracting the effects of gravity. Embedded bioprinting facilitates the production of scalable, vascularized tissues *in vitro* by allowing the direct bioprinting of perfusable vascular-like networks ^[8]^.

In an early contribution to the field, omnidirectional printing of 3D biomimetic microvascular networks was reported, which involved the printing of fugitive ink filaments within a photo-curable gel reservoir. This gel reservoir served as both a physical support for the patterned features and a means to facilitate genuine omnidirectional freeform fabrication ^[9]^. A few years later, the development of granular gels that can fluidize at the point of injection and then rapidly solidify, trapping injected material in place was reported ^[10]^. This method of creating 3D structures reduces the influence of surface tension, gravity, and particle diffusion, thus enabling the use of an extensive range of materials. Silicones, hydrogels, colloids, and living cells were used to fabricate intricate 3D structures ^[10]^. Concurrently, a technique known as Freeform Reversible Embedding of Suspended Hydrogels (FRESH) was introduced. This approach used a thermoreversible support bath, facilitating the deposition of hydrogels in intricate 3D biological structures within a secondary hydrogel support bath, comprised of gelatin microparticles. Upon successfully printing functional hydrogel inks, the sacrificial ink was dissolved, reclaiming bioprinted structures thus allowing the fabrication of complex constructs, including vasculature and organ models ^[11]^. Recently, Fang et al. reported a bioprinting strategy termed sequential printing in a reversible ink template (SPIRIT), which utilizes a microgel-based biphasic bioink that serves dual roles as both a bioink and a suspension medium. This capability is facilitated by the bioink’s shear-thinning and self-healing characteristics, allowing for embedded bioprinting. Notably, this strategy proved successful in the fabrication of a ventricle model with an intricate, perfusable vascular network_[12]._

While these methods can produce vascularized tissues with nearly physiological cell density, they face limitations in replicating the external and internal geometries of native tissues simultaneously. The fundamental problem of bioprinting complex internal vasculature all at once lies in the intricate nature of blood vessels, as they require precise spatial arrangements and interconnected networks. Attempting to print them all at once can lead to challenges in accurately recreating the intricate structures, potentially resulting in functional issues. Embedded bioprinting addresses this problem by focusing on bioprinting a scaffold first and then infusing it with cells and a biocompatible material to stimulate vascular growth. This approach allows for more control over vascular development and ensures better integration with the surrounding tissue, increasing the chances of successful bioprinting of complex vasculature. Additionally, there is a demand for the creation of a hydrogel that can serve dual roles as a bioink and a suspension medium, particularly for embedded bioprinting purposes.

Drawing from these concepts, we present a novel advancement in embedded bioprinting, henceforth referred to as “intra-embedded bioprinting (IEB).” Akin to nesting a Matryoshka doll, the act of embedded bioprinting is executed within a larger, pre-existing embedded bioprinted structures. This was possible as the proposed support material can be used as an extrudable bioink as well as a support bath for the follow-up (i) bioinks to deliver viable cells, or (ii) sacrificial inks to create perfusable vasculature. Herein, first, an extrudable ink was bioprinted within a sacrificial or functional support bath to mimic the overall 3D shape of the targeted tissue (**Figure 1**). Next, bioinks, including cell-laden hydrogels or spheroids, were deposited within the previously embedded bioprinted constructs. Also, a sacrificial ink was bioprinted within the bioprinted structures, which upon removal yields perfusable channels inside a complex-shaped tissue construct. To showcase the capabilities of IEB, a miniaturized pancreatic model was 3D printed towards organ modeling using a photo-crosslinkable silicone composite support bath followed by the development of nested structures, and the fabrication and validation of a breast tumor model. Thus, IEB was proposed here as an advanced version of embedded bioprinting and has the potential to develop heterotypic tissues or tissue models by converging multiple technologies such as extrusion-based bioprinting (EBB) and aspiration-assisted bioprinting (AAB).

**Figure 1.**
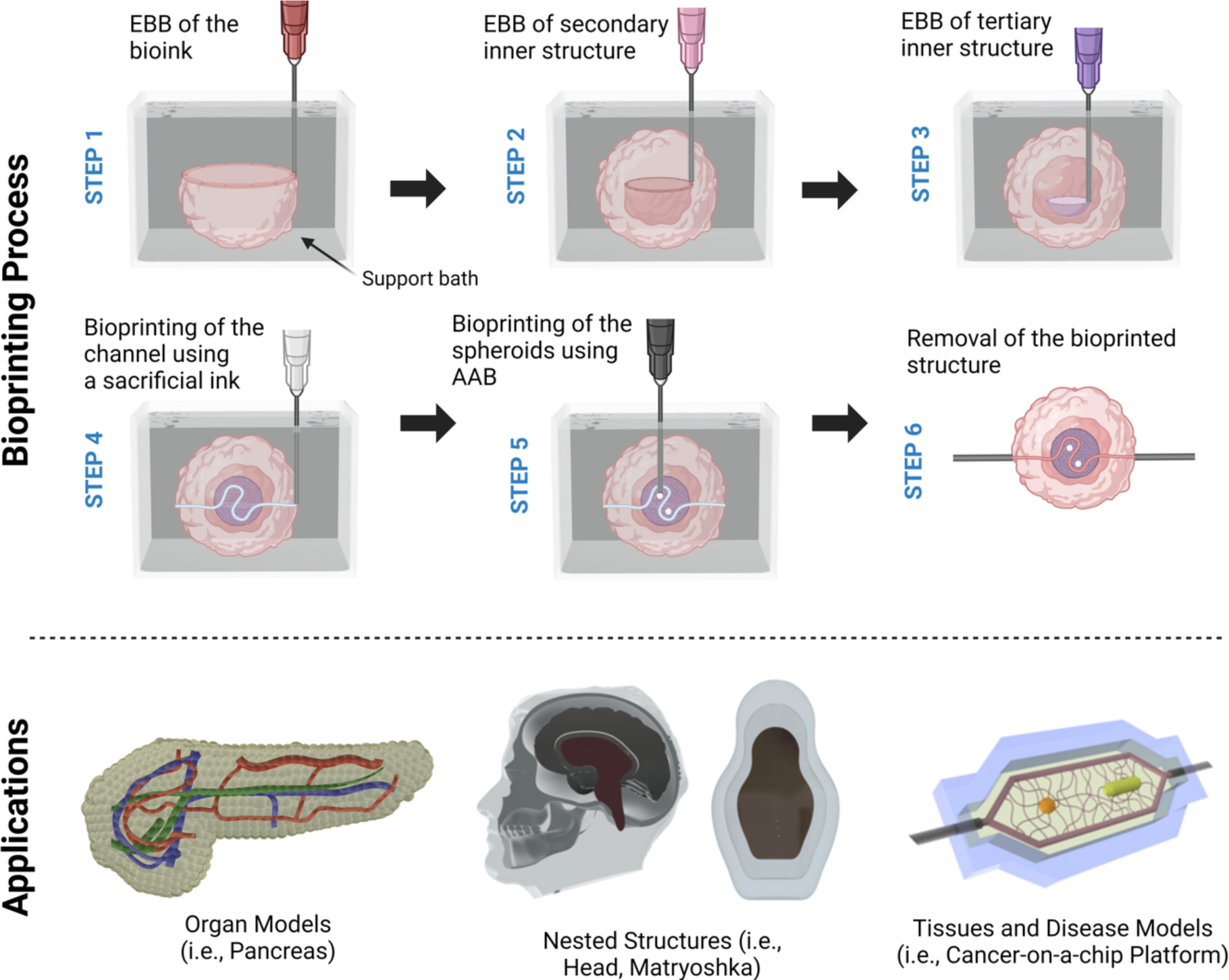
A schematic illustrating the step-by-step process of IEB and its applications. (created with Biorender.com).

## 2. Results and Discussion

In this study, IEB was proposed as an advanced version of embedded bioprinting to fabricate complex-shaped organ models, tissues, or cancer-on-a-chip platforms with intricate anatomy or internal structures. With this method, we demonstrated the capability to bioprint inside previously bioprinted constructs. To achieve IEB, first, an extrudable ink was bioprinted inside a support bath following the conventional embedded bioprinting approach. Then, the bioprinted structure was used as a functional support bath to bioprint the subsequent inks. In this manner, we demonstrated that the extruded inks could also be used as support baths. The subsequent inks could be sacrificial inks to generate perfusable channels, bioinks to bioprint viable structures, spheroids, or organoids to develop heterotypic complex tissues (**Figure 1**).

### 2.1. Hydrogel Synthesis and Analysis

Xanthan gum (XaG) and gelatin were methacrylated with methacrylic anhydride to form methacrylated xanthan gum (XaGMA) and methacrylated gelatin (GelMA), respectively, and 0.2% (w/v) Lithium phenyl (2,4,6-trimethylbenzoyl) phosphinate (LAP) was used as a photocrosslinker. The mechanisms for both reactions are presented in **Figure 2A**. The methacrylation reaction was validated with fourier-transform infrared spectroscopy (FTIR) and ^1^H nuclear magnetic resonance (NMR) tests. FTIR data of GelMA and XaGMA suggested that the peak changes at 1640–1710 cm^−1^ range were due to C=O and C=C bonds in methacrylic anhydride ^[13–15]^ (**Figures 2B1-B2**). NMR data suggested that the peak at 2.9 ppm represented the methylene of lysine amino acid and its reduction from gelatin to GelMA signified the reduction in the associated proton intensity due to the modification done by methacrylic anhydride. Further, the appearance of the peaks at 5.4 and 5.7 ppm in GelMA as well as in XaGMA denotes the protons of methacrylate vinyl group of methacrylic anhydride ^[16]^ (**Figures 2B3-B4**). To investigate the ultra-morphology of the crosslinked hydrogels, scanning electron microscope (SEM) imaging was also performed (**Figure 2C**). The samples exhibited pore sizes, typically in the range of approximately 20 to 100 µm, which allowed for the migration of cells (**Figure S1**). The pore distribution revealed that, in comparison, XaGMA samples exhibited larger pores, while GelMA had smaller pores with the XaGMA/GelMA composite pore size falling in between. This pore size range can be considered to be suitable for facilitating cell migration and promoting their proliferation ^[17]^

**Figure 2.**
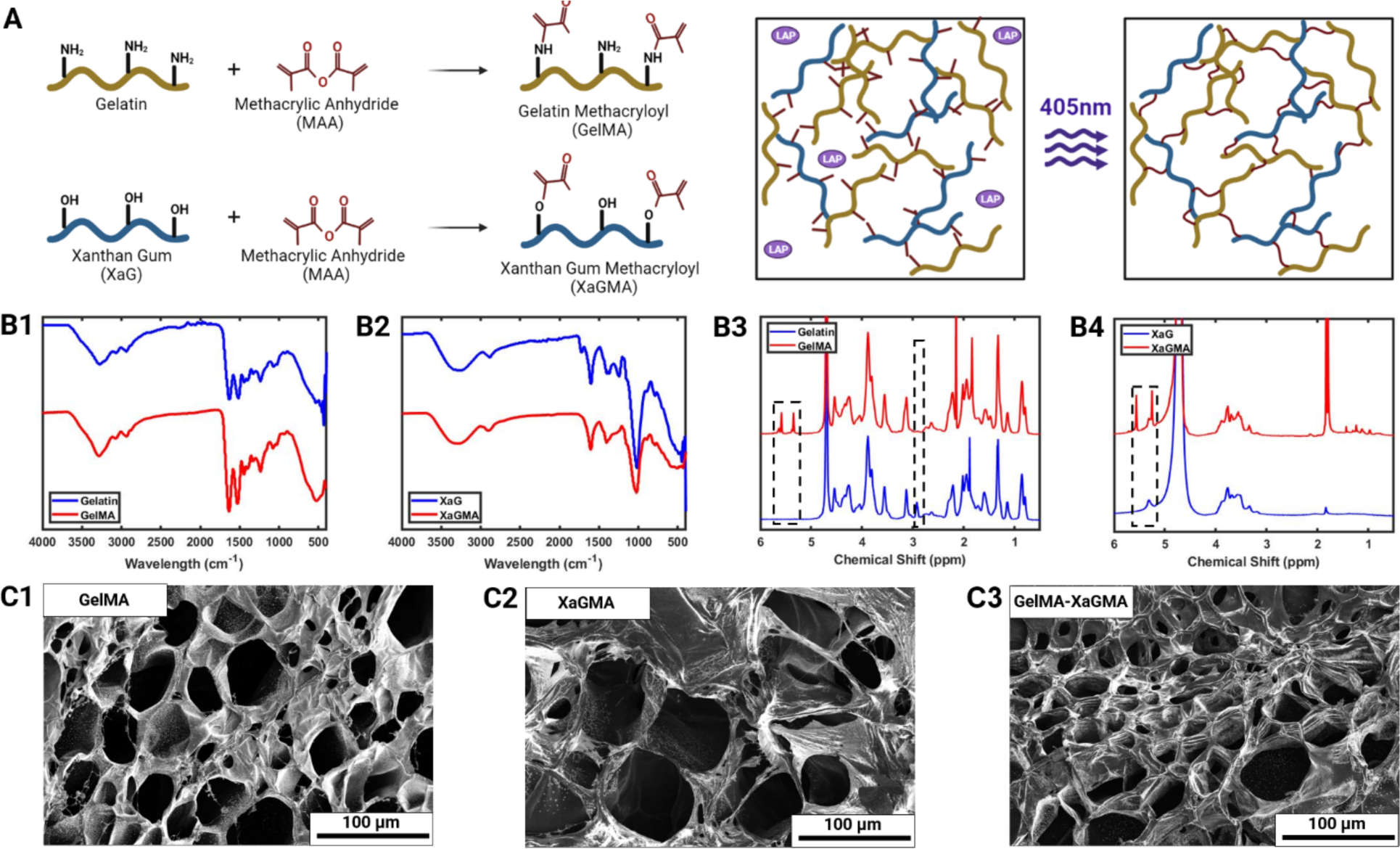
XaGMA and GelMA hydrogels. (A) Synthesis and crosslinking of XaGMA and GelMA. FTIR of (B1) GelMA and (B2) XaGMA. ^1^H NMR of (B3) GelMA and (B4) XaGMA. SEM images of (C1) GelMA, (C2) XaGMA, and (C3) the XaGMA/GelMA composite.

### 2.2. Assessment of Printability

In this study, analysis for rheological properties was performed to understand the flow and deformation mechanisms of the prepared composite hydrogels. The rheological findings explained the extrusion, shear-thinning, and recovery characteristics of these hydrogels which were crucial to examine for their compatibility to be used as an extrudable ink and a support bath. The results indicated that the methacrylation of XaG to XaGMA did not change its rheological properties as the viscosity, shear thinning, and recovery were similar for XaG and XaGMA for a given concentration (**Figure S2**). The amplitude sweep test depicted that storage modulus (G’) > loss modulus (G”) at lower shear strain, indicating that the material behaved like a solid, while G’ < G” at higher strain values suggesting that the material had fluid-like properties (**Figure 3A**). In the elastic region, the storage modulus of 1, 1.5, and 2% XaGMA was found to be ∼35, 60, and 120 Pa, respectively, which was unaffected by the addition of GelMA. The transition from solid to fluid, which is called the yield point, occurred at a shear strain of ∼140, 120, and 115% with the yield stress of ∼20, 35, and 65 Pa for 1, 1.5, and 2% XaGMA, respectively (**Figure S3A**). The yield stress in this range was reported as suitable for EBB as an ink as well as a support bath. The frequency sweep test was performed within the viscoelastic region (shear strain 0.1%), and it was found that G’ > G” indicated the material was elastic for the whole range of angular frequency from 0.1 to 100 rad/s (**Figure S3B**). The flow rate sweep test showed that the viscosity decreased with an increase in shear rate for all the samples, which validated that the hydrogels were shear thinning (**Figure S3C**). Further, the viscosities of XaGMA and XaGMA/GelMA were found similar for the same XaGMA concentration, and they increased only with the increase in the concentration of XaGMA, which corroborates with the above-mentioned rheological results. The thixotropy or self-healing tests were carried out at a cyclic high shear rate (100 s^−1^) and low shear rate (1 s^−1^) (**Figure 3B**), showing that the viscosity of all the samples quickly recovered, suggesting that the materials had a quick recovery time, making them ideal for use as an extrudable ink as well as a support bath.

**Figure 3.**
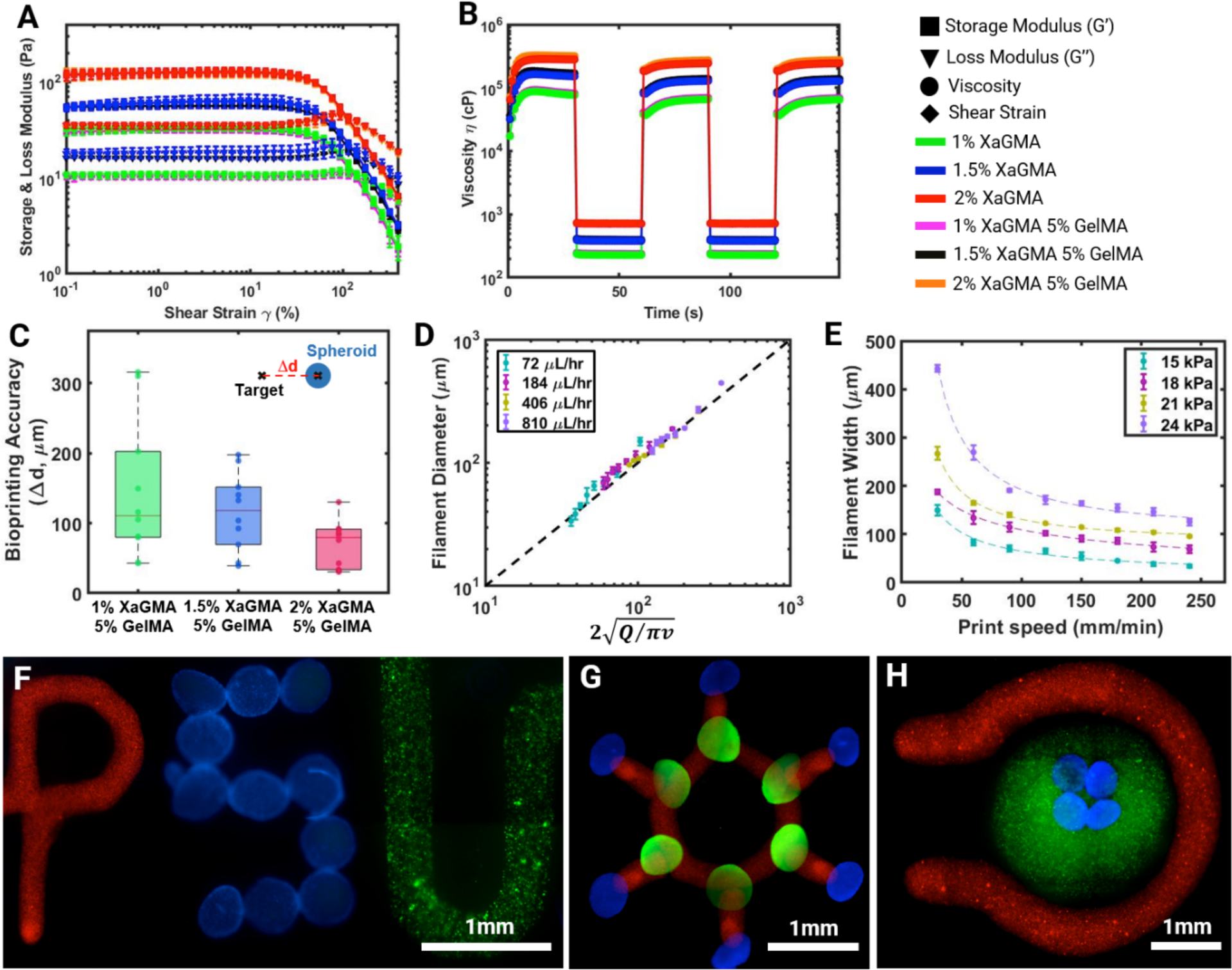
Assessment of printability and IEB demonstrations. (A) Amplitude sweep test to measure the storage modulus, loss modulus, and yield stress at a strain ranging from 0.1 to 300%, and (B) self-healing thixotropy test to validate the material recovery behavior under cyclic high (100 s^−1^) and low (1 s^−1^) shear rate (*n*=3). (C) Bioprinting accuracy of spheroids (*n*=10), (D) filament width of sacrificial inks in line with the fluid continuity equation, where *Q* and *v* represent the flow rate and nozzle speed, respectively (*n*=6), and (E) filament diameter versus printing speed curves at 15, 81, 21, and 24 kPa pressure levels (*n*=6). (F) Printing demonstration of PSU design, where ‘P’ was printed with the XaG sacrificial ink (red), ‘S’ with DAPI-stained hMSC spheroids (Blue), and ‘U’ with XaG bioink containing GFP^+^ MDA-MB-231 cells (green), (G) printing demonstration of a benzene design with XaG ink (red) and hMSC spheroids (blue and green), and (H) IEB of DAPI-stained hMSC spheroids (blue) inside 1.5% XaG loaded with GFP^+^ MDA-MB-231 cells (green) and the XaG ink (red) around it.

To test the suitability of XaG as a potential support bath, EBB and AAB were performed in tandem in XaG-based support bath. For EBB, XaG itself was used as the ink while hMSC spheroids were bioprinted via AAB. For the initial testing, filaments ranging from 100 to 600 µm and multiple spheroids were bioprinted (**Figures S4**). The accuracy of AAB was measured, and it was found that the accuracy, which was reflected as the distance between the target location and the bioprinted spheroid location, was within 300 µm and improved with increasing XaGMA concentrations (**Figure 3C**). This might be attributed to the high viscosity of XaG, which formed a stable bath and entraped spheroids in their originally bioprinted locations thus mitigating their movements post bioprinting.

Further, the printability of the sacrificial ink (1.5% XaG) was characterized by measuring the size (diameter) of printed filaments at different printing speeds and volumetric flow rates. Sacrificial inks were extruded using blunt straight needles (32G, 100 µm inner and 240 µm outer diameter). By altering the above-mentioned parameters, filament diameter could be adjusted. The measured filament diameters were compared with the theoretical values calculated from the fluid continuity equation ^[18]^. This equation takes parameters as volumetric flow rate (*Q*), nozzle speed (*v*) and predicts the diameter of the produced filament (*d*) by following the relationship *π(d/2)^2^=Q/v*. It was found that at different combinations of material flow rate and nozzle speed, the experimental filament width matched with the predicted diameter calculated from the fluid continuity equation (**Figure 3D**). It was also observed that the filament diameter decreased as the printing speed increased or the flow rate decreased (**Figure 3E**). Overall, the results indicate that the filament diameter could be closely predicted by following the flow continuity equation.

To demonstrate the versatility of the presented technique, the sacrificial ink loaded with red fluorescent particles, diamidino-2-phenylindole (DAPI)-stained human mesenchymal stem cells (hMSC) spheroids and the bioink (1.5% XaG)-laden green fluorescent protein (GFP)^+^ MDA-MB-231 cells were bioprinted in the shape of the abbreviation for Penn State University (PSU) (**Figure 3F**). Also, a benzene design was made with sacrificial ink loaded with red fluorescent particles, and DAPI- and 488 phalloidin-stained hMSC spheroids (**Figure 3G**). Finally, DAPI-stained spheroids were bioprinted inside the bioink (1.5% XaG) loaded with GFP^+^ MDA-MB-231 cells (**Figure 3H**).

The above-mentioned printing tests validated that the support bath was compatible with both EBB and AAB. The integration of EBB and AAB thus allowed the development of heterotypic tissue models that contained spheroids placed with AAB as well as bioprinted filaments or sacrificial filaments using EBB. Therefore, it opens opportunities for developing complex functional organ models by converging multiple bioprinting technologies.

### 2.3. Assessment of cytocompatibility

During the initial optimization of the XaGMA/GelMA composite, we conducted biocompatibility experiments using 3T3 and MDA-MB-231 cells. The cells were incorporated into composite hydrogels with varying concentrations of XaG, specifically 1% XaG/5% GelMA, 1.5% XaG/5% GelMA, and 2% XaG/5% GelMA and assessed using the LIVE/DEAD staining. The viability of both cell types across all samples exceeded 90% for a duration of up to 7 days (**Figure S5**). Accordingly, based on the initial printing characterization and cell viability results, only the 1.5% XaG/5% GelMA composite was considered for further studies. Thus, adipose-derived stem cells (ADSCs) (being stem cells and used in the current study to develop cancer-on-a-chip model) were used to assess the biocompatibility of 1.5% XaG/5% GelMA composite hydrogels over 14 days, both in the cast and bioprinted configurations. The results demonstrated >90% cell viability at Day 7 which was maintained at Day 14 in the bioprinted constructs (**Figure 4**). Additionally, cell spreading could be observed in bioprinted constructs within 7 days, suggesting that the bioprinted constructs provided a conducive environment for cellular growth. Further, ADSCs maintained their viablilty on cast samples as well; however, limited spreading was observed by Day 14, which might be due to their bulk hydrogel nature as opposed to the bioprinted filaments (**Figure S6**). Overall, these findings emphasize on the high biocompatibility profile of the XaG/GelMA composites across diverse cell types and concentrations, indicating their potential for biomedical and tissue engineering applications.

**Figure 4.**
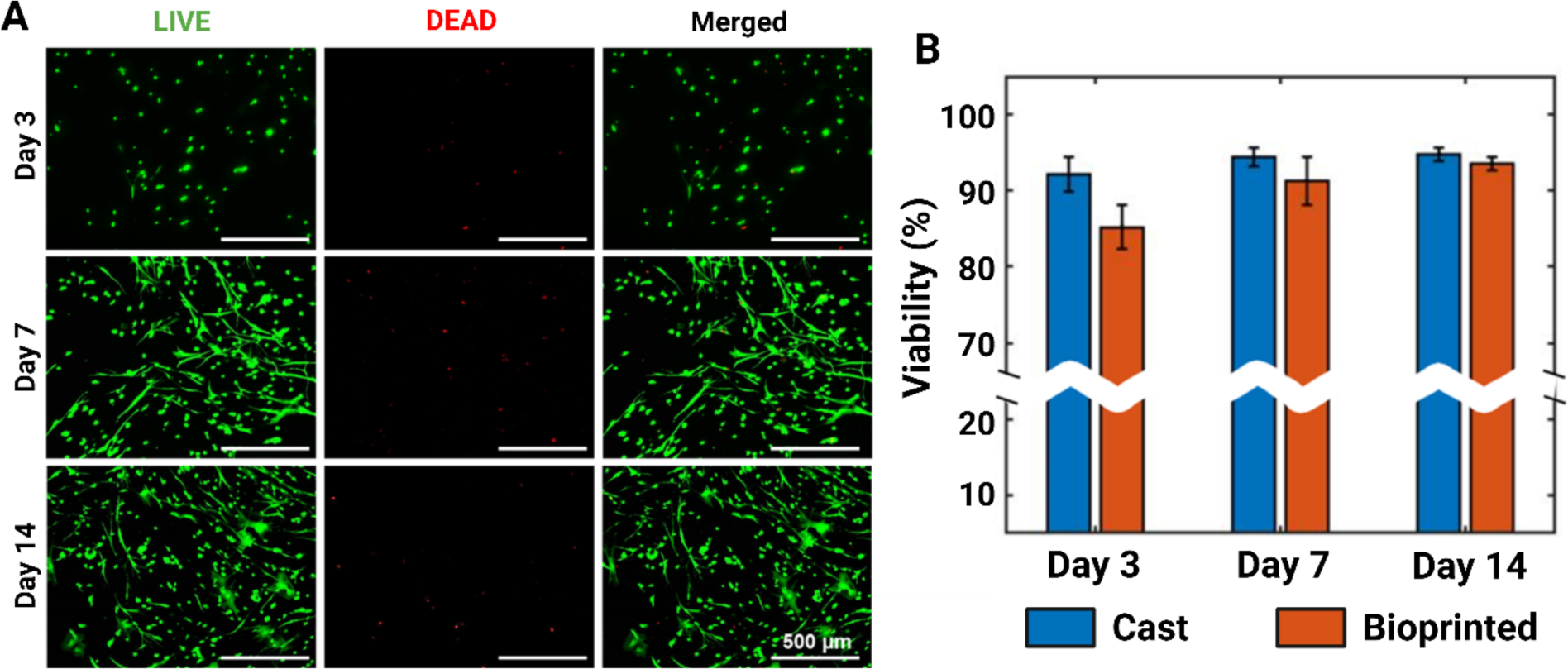
Viability of cells in 1.5% XaG/5% GelMA. (A) Representative LIVE/DEAD images of bioprinted ADSCs on Days 3, 7, and 14. (B) Viability of ADSCs on cast and bioprinted constructs on Day 3, 7 and 14 (*n* = 3).

### 2.4. Exemplification of IEB

Having established the capability of IEB as demonstrated in **Figures 3F-H**, we then aimed to fabricate a scaled-down model (18 x 9 x 6 mm; length x height x depth) of a human pancreas with associated vasculature and bile ducts. We started with designing the model including arteries and veins that support the pancreas and pancreatic ducts (**Figure 5A1**). The model was fabricated using a photo-crosslinkable silicone composite in the XaG-based support bath, where the vasculature was printed using oil-based dye-containing XaG (**Figure 5A2, Supporting Movie 1**). The printed model was transparent allowing for real-time monitoring of the vasculature printing. After 3D printing, the dimensions were comparable to the original file (**Figure 5A3**). Previously, a full-size model of an adult human heart made of alginate was reported ^[19]^. While we have presented a miniaturized pancreas model, the method presented here is extensible to printing a full-size organ model as well as other organ models (e.g., liver, heart) at a realistic size. Such organ models hold a significant role in surgical training procedures, as well as in non-surgical contexts such as communication and education. To further, exemplify the IEB method, another organ, a head phantom with anatomical structures including skull, cerebral cortex, and brain stem, was 3D printed inside the support bath (**Figures 5B1-B2, Supporting Movie 2**). Additionally, a Matryoshka doll was printed using four precisely crafted layers, exemplifying the method’s significant capability to manufacture complex and nested structures (**Figures 5C1-C3, Supporting Movie 3**). Each layer was precisely encapsulated into the previously printed one, allowing for the creation of intricate and precise geometries. This level of control over layering is crucial in various technical applications, including microfluidic device production, where intricate nested structures are crucial for optimal performance and functionality ^[20]^.

**Figure 5.**
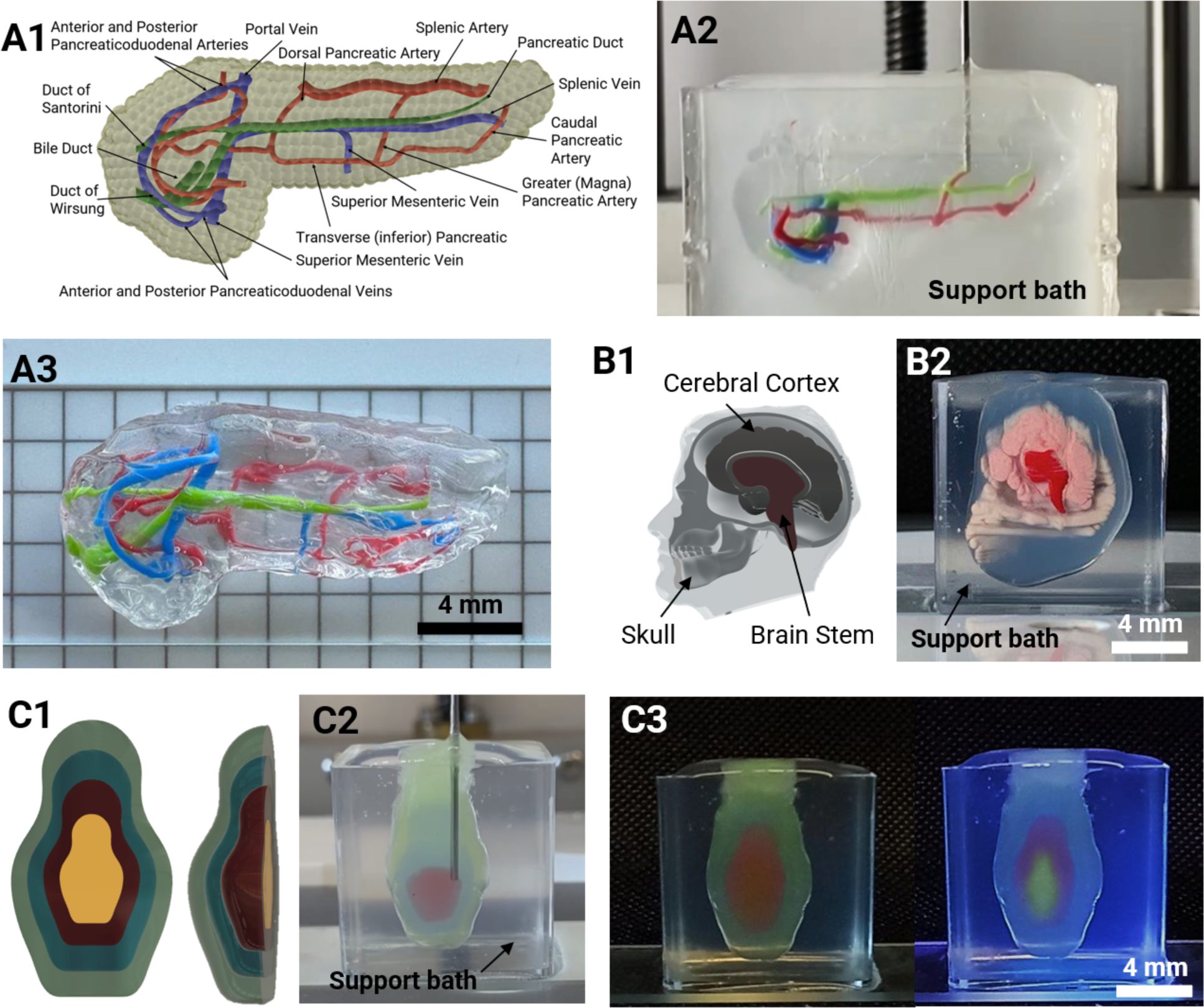
Printing nested structures. (A1) Desing of a pancreatic model with the vasculature and bile ducts, (A2) 3D printing in action, and (A3) the 3D printed model. (B1) The design of a head phantom with anatomical structures including skull, cerebral cortex, and brain stem, and (B2) the corresponding four-nested (labelled 1-4) 3D printed structure. (C1) The design of a Matryoshka doll, (C2) the print in progress, and (C3) the corresponding four-nested (labelled 1-4) 3D printed structure.

Towards microfluidics and cancer-on-a-chip application, an *in-vitro* breast tumor model was fabricated using IEB (**Figures 6A-B**). The model consisted of a XaGMA/GelMA support bath into which a bioink composed of XaGMA/GelMA-laden adipose cells were deposited using EBB in a spherical shape. The adipose cells were obtained from ADSCs, and their differentiation was confirmed using the adipogenic markers including Peroxisome proliferator-activated receptor gamma (PPAR-G), the adipocyte fatty acid-binding protein (aP2), and Adiponectin (ADIPOQ). The expression of these adipogenic genes was significantly higher on Day 15 as compared to ADSCs cultured with basal medium (**Figure S7**). The adipose cells in the bioink provided a breast tissue-like environment ^[21]^. Next, a sacrificial ink composed of XaG was printed across the support bath passing through the bioprinted breast model, forming a hollow channel upon removal of the sacrificial ink. Finally, into the model and close to the channel (200 to 400 µm), tumor spheroids (composed of MDA-MB-231 and ADSCs) were bioprinted using AAB (**Figures 6C-E, Supporting Movie 4**). Based on a previous report ^[22]^, we here used a mixture of the two cell types to form stable spheroids as MDA-MB-231 spheroids were structurally weak. Additionally, ADSCs are one of the major stromal cells in the breast cancer microenvironment that promote cancer progression ^[23]^. The channel of the developed model was then perfused with Doxorubicin (DOX) and its effect on tumor spheroids was assessed. The anthracycline DOX is a common first-line therapy for numerous human pathological conditions, such as breast cancer ^[24]^. Being a well-established drug, it was used in our study to validate the functionality of the model. The results showed that the measured volume of the tumor spheroids reduced significantly (p≤0.05) on Day 3 of treatment with DOX (**Figures 6F-G**). The results were in agreement with Wang et al., where a 3D bioprinted breast cancer model consisting of 21PT breast tumor cells line and adipose derived-MSC (adMSC) was developed using dual hydrogel-based bioinks ^[25]^. They reported that the cleaved Caspase-3 positive cell percentage was significantly lower in the bioprinted constructs with adMSC and 21PT than that in the cancer cell alone constructs, in response to a low DOX dose (i.e., 1 and 10 µg/mL). The results validate that the developed IEB method can be used to fabricate cancer-on-a-chip models for testing anticancer drugs and other therapeutics including cancer immunotherapy.

**Figure 6.**
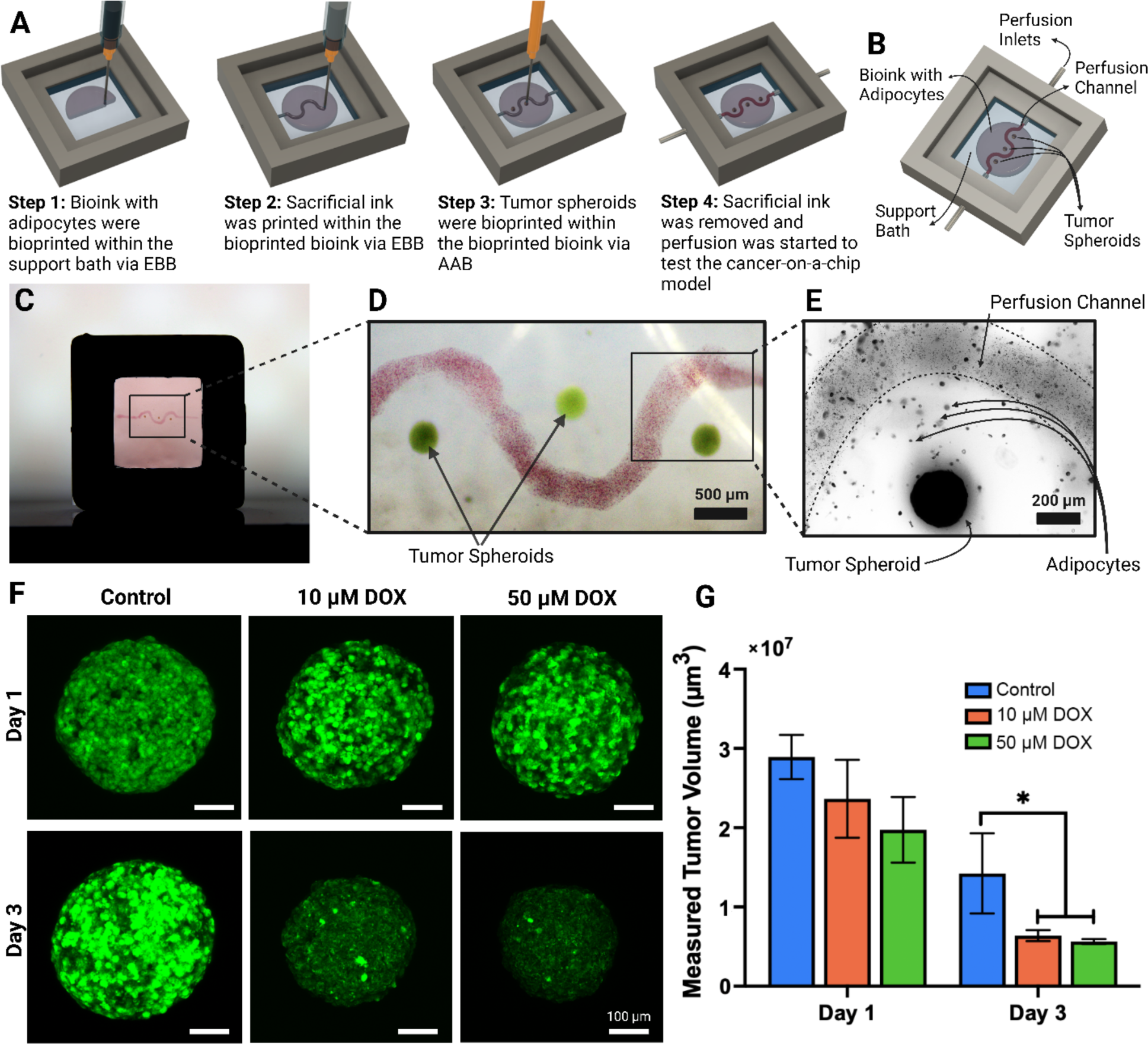
Cancer-on-a-chip platform. (A) The schematic of the developed cancer-on-a-chip model, (B) the components and set-up of the device, (C) the bioprinted breast tumor model with (D) a magnified image showing the channel and tumors, and (E) adipocytes within the bioink. (F) Tumor spheroids with 10 and 50 µM DOX perfusion at Days 1 and 3 (spheroids without DOX treatment were used as a control), and (G) change in tumor volume post treatment with DOX (*n* = 3; *p ≤ 0.05).

## 3. Conclusion

In this paper, with inspiration from the Matryoshka doll, we introduced an innovative method called intra-embedded bioprinting as an advancement over existing embedded printing methods. Utilizing XaG- and GelMA-based inks, we demonstrated their dual purpose use as a bioink and support bath. To illustrate the potential of IEB, we created a scaled-down model of pancreas that closely matched the original design, affirming the precision of IEB. Additionally, a 4-layered head phantom and Matryoshka doll were 3D printed, exemplifying IEB’s capabilities to manufacture complex and nested structures. Furthermore, using innovative coupling between EBB and AAB, we successfully bioprinted a cancer-on-a-chip model and validated its response to chemotherapeutics. While we demonstrated IEB’s feasibility using only a limited number of biomaterials, it is logical to infer its applicability to other gels with yield stress and self-recovery properties. Particularly, IEB brings forth an efficient approach for fabricating complex tissue constructs with embedded internal features, such as freeform vascular networks, expediting the engineering of complex tissue and organ structures, opening new possibilities in the fields of biofabrication and therapeutic advancements.

## 4. Experimental Section

### Materials

Xanthan Gum from *Xanthomonas campestris*, gelatin from cold water fish skin, methacrylate anhydride (MAA), sodium carbonate (Na_2_CO_3_), sodium hydroxide (NaOH), sodium alginate, and 2,2-Dimethoxy-2-phenylacetophenone were acquired from Sigma-Aldrich (St. Louis, MO, USA). LAP was acquired from TCI chemicals (Tokyo, Japan). 12-14 kDa molecular weight cut-off (MWCO) dialysis membrane was acquired from Repligen (Waltham, MA, USA). Dulbecco’s phosphate-buffered saline (DPBS) was acquired from Corning (NY, USA). Vinyl terminated (4-6% diphenylsiloxane) dimethylsiloxane copolymer (PDV-0525) and (mercaptopropyl 13-17%) methylsiloxane - dimethylsiloxane copolymer (SMS-142) were acquired from Gelest (Morrisville, PA, USA). HMDS-treated silica (AEROSIL R 812 S) was acquired from Evonik (Essen, Germany). Thixotropic agent THI-VEX™ (ThA) was obtained from Smooth-On (Macungie, PA, USA).

### Preparation of XaGMA and GelMA

To obtain XaGMA, we have used a modified version of a previously reported method ^[26]^. Briefly, 7.5 g of xanthan gum (XaG) was blended in 300 mL of 0.25 M carbonate-bicarbonate buffer (CB) at pH 9 for 4 min using a commercial blender (Magic Bullet, CA, USA). 400 µL of MAA was added in 4 intervals (100 µL in each interval). At each interval, MAA was blended for 4 min and rested for 30 min before the next interval. The final blend was left to react overnight at 4 °C and dialyzed with a 12-14 kDa MWCO dialysis membrane at 40 °C for four days. Following dialysis, the solution was lyophilized in a freeze drier (FreeZone 18 L Console Freeze Dry System, Labconco, Kansas City, MO, USA) and stored at 4 °C for further use.

To obtain GelMA, gelatin was methacrylated using a previously reported method ^[27,28]^. Briefly, 0.1 M CB solution was prepared and pH was adjusted to 9 using NaOH. Then, 20 g of gelatin was dissolved in 100 mL CB at 50 ⁰ C using a magnetic stirrer. Once gelatin was completely dissolved, pH was again adjusted to 9 using NaOH. After the adjustment, 500 µL of MAA was added in 5 intervals (100 µL of MAA was added for each interval of 15 min, and pH was adjusted to 9 at each addition). After the addition of MAA, the reaction was maintained for 3 h at 55 °C. Then, the reaction was quenched by pH to 7.4. Next, the solution was filtered and kept under dialysis with a 12-14 kDa MWCO dialysis membrane for four days at room temperature. After the dialysis, the solution was lyophilized using a freeze drier and stored at 4 ⁰ C for further use.

### Hydrogel Preparation

A stock solution of 20% (w/v) GelMA was prepared by dissolving 20 g of GelMA in 100 mL DPBS on a magnetic stirrer at 60 °C. After dissolving, the solution was sterilized with a 0.22 µm vacuum filter. Stock solution of 3% (w/v) XaG and XaGMA was prepared by blending 9 g of each XaG and XaGMA separately in 300 mL DPBS for 5 min. After blending, the solution was sterilized by autoclaving. A stock solution of 2% (w/v) LAP was prepared by dissolving 20 mg of LAP in 1 mL DPBS and sterilized with a 0.22 µm syringe filter. For all samples, the final LAP concentration was 0.2% (w/v). To prepare the mixtures, all solutions (GelMA, DPBS or cell media, LAP, XaG, or XaGMA) were transferred to 2 mL centrifuge tubes and vortexed for 5 min. After vortexing, the mixtures were placed into an ultrasonic sonicator for 2.5 min and centrifuged for 5 min at 4,000 rpm to remove air bubbles. After casting and bioprinting processes were completed, hydrogels were crosslinked by exposing them to a 358 mW/cm^2^, 405nm light source (CR-UV light, Creality, Shenzhen, China) for 1 min.

### Fourier-transform infrared spectroscopy

FTIR spectra of powder form of gelatin, GelMA, XaG and XaGMA were recorded with a spectral range from 3,500 to 400 cm^−1^ and 16 scans on an attenuated total reflection FTIR spectrometer (Nicolet 6700, Thermo Fisher Scientific, MA).

### Nuclear Magnetic Resonance Spectroscopy

Gelatin, GelMA, XaG and XaGMA powder were dissolved in deuterium oxide (D_2_O) to obtain 20 mg/mL solutions. Further, proton NMR spectra were obtained using a Bruker AVIII-HD-500 (Billerica, MA, USA) spectrometer with a Bruker Ascend magnet of 500 MHz. The degree of functionalization was calculated using NMR and it was defined as the number of methacrylated groups attached to gelatin divided by the total number of amine groups of the unfunctionalized gelatin ^[29]^.

### Field Emission Scanning Electron Microscopy

A field emission scanning electron microscope (FESEM) (Apreo SEM, Waltham, MA, USA) was used to demonstrate the surface topography and pores of crosslinked samples. Samples were placed at −80 °C overnight, then lyophilized in a freeze dryer. Freeze-dried samples were placed on a carbon tape and were sputter coated with a 7.5 nm iridium layer using a Leica EM ACE600 sputter coater (Wetzlar, Germany). After coating, the samples were imaged at an accelerating voltage of 3 kV and beam current of 6.3 pA. The pore sizes were measured from the SEM micrographs using ImageJ software. Three images were taken for each sample. All visible pores in images were considered and a graph demonstrating the pore size distribution was plotted.

### Rheological Characterization

Rheological measurements were performed on an MCR 302 rheometer (Anton Paar, Graz, Austria) equipped with a 25-mm parallel plate at 23 °C as per published methods ^[30,31]^. An amplitude sweep test was performed within a range of 0.1 to 300% shear strain at a constant frequency of 1 Hz. A frequency sweep test was performed within a range of 0.1 to 100 rad s^−1^ angular frequency at a constant shear strain of 0.1%. A shear rate sweep test was performed within a range of 0.1 to 100 s^−1^ shear rate at a constant frequency of 1 Hz. Self-recovering thixotropy test was performed at a cyclic low shear rate of 0.1 s^−1^ for 1 min followed by a high shear rate of 100 s^−1^ for 1 min. The cycle was repeated three times and the recovery of viscosity was measured.

### Cell culture

GFP^+^ MDA-MB-231 metastatic breast cancer cells (passage 25-30) were cultured in Roswell Park Memorial Institute (RPMI) 1640 media supplemented with 10% (v/v) fetal bovine serum (FBS) (Life Technologies, Grand Island, NY, USA), 1 mM Glutamine (Life Technologies, Carlsbad, CA, USA) and 1% (v/v) penicillin-streptomycin (PS, Life Technologies). Mouse embryonic fibroblasts (3T3 Cells, passage 32-35) were obtained from American Type Culture Collection (ATCC, Manassas, VA, USA) and cultured in a complete growth medium composed of 1% (v/v) PS, 10% (v/v) FBS, and 89% (v/v) Dulbecco’s Modified Eagle Medium (DMEM). Human mesenchymal stem cells (Passage 5-7) (commercially obtained from Rooster Bio (Frederick, MD)) were cultured in hMSC growth media (R&D Systems, Minneapolis, MN, USA). Adipose-derived stem cells (Passage 3-5) (Lonza, Walkersville, MD, USA) were cultured in a basal medium consisting of DMEM/F12, supplemented with 20% (v/v) FBS (R&D Systems), 1% (v/v) PS at 37 °C with 5% CO_2_. Cell media was changed every other day.

### Spheroid Fabrication

Breast tumor spheroids were fabricated using the method described before ^[32]^. 2 × 10^3^ MDA-MB-231 breast cancer cells were mixed with 2 × 10^3^ ADSCs and 200 µL of this cell suspension was seeded into each well of a U-bottom 96-well plate with cell-repellent surface for 24 h (Greiner bio-one, Frickenhausen, Germany) to form the spheroids. They were maintained in ADSC growth media to allow compaction at 37 °C. Similarly, hMSCs spheroids were formed using ∼1 × 10^6^ cells per mL into each well of a U-bottom 96-well plate with cell-repellent surface and incubated for 24 h.

### Adipogenic Differentiation and Validation

ADSCs were cultured in basal medium. When they reached 80% confluence, they were treated with a human adipocyte differentiation medium (Cell Applications, Inc., San Diego, CA, USA) for 15 days. To confirm the differentiation of ADSCs into adipogenic cells, gene expression was performed. ADSCs were cultured and differentiated on a 175 cm^3^-cell culture flasks. After 15 days, cells were washed twice with PBS, detached with Trypsin and collected by centrifugation at 1,600 xg for 4 min and homogenized in TRIzol (Life Technologies). PureLink RNA Mini Kit (Thermo Fisher) was used to isolate total RNA from cells according to the manufacturer’s protocol. A Nanodrop (Thermo Fisher) was used for measuring the RNA concentration, and reverse transcription was performed using AccuPower® CycleScript RT PreMix (BIONEER, Daejeon, Korea) following the manufacturer’s instructions. Gene expression was performed quantitatively with SYBR Green (Thermo Fisher) using a QuantStudio 3 PCR system (Thermo Fisher). Adipogenic genes tested included Peroxisome proliferator-activated receptor gamma (PPAR-G), the adipocyte fatty acid-binding protein (aP2), and Adiponectin (ADIPOQ) (see **Table S1** for the list of primers). All genes were normalized to the expression of a β-actin gene, and the 2^−ΔΔCT^ method was used to calculate gene expression levels relative to Day 1. ADSCs cultured with basal medium were used as a control.

### Cell Viability Assessment

To measure the cell viability, 100 µL composite hydrogel of 1, 1.5, or 2% XaGMA mixed with 5% GelMA containing 1 x 10^5^ ADSCs were cast into 24-well plates and crosslinked by exposing them to a 405-nm light source for 1 min. Cell viability was measured on Days 1, 7, and 14 using the LIVE/DEAD assay. For the viability assessment, cells embedded in hydrogels were rinsed with DPBS three times, followed by staining with calcein AM (2 µM) and ethidium homodimer-1 (EthD-1) (4 µM) solutions for 1 h in an incubator. After incubation, hydrogels were rinsed with DPBS three times and imaged using a Zeiss Axio Observer microscope (Zeiss, Jena, Germany). The cell viability was quantified by thresholding the obtained microscopic images using ImageJ and the area covered by live and dead cells was calculated. During the initial optimization and biocompatibility testing of hydrogel samples, 3T3 and MDA-MB-231 cells were used, and the viability of cells was estimated using the LIVE/DEAD assay.

### Extrusion-based Bioprinting and its Characterization

For EBB during IEB, the prepared composite hydrogel (1.5% XaGMA/ 5% GelMA) was bioprinted inside the support bath (1.5% XaG) using a 3D bioprinter (Inkredible+, Cellink, Gothenburg, Sweden). After bioprinting the first layer, the sacrificial ink (1.5% XaG) was printed using a 32G needle (100 µm inner and 240 µm outside diameter) within the initially bioprinted composite hydrogel for characterization purposes. To explore the effect of printing speed and applied pressure on the filament size, the printing speed was gradually increased from 30 to 240 mm/min at extrusion pressures of 15, 18, 21, and 24 kPa. Also, flow rates for the applied pressures were calculated by extruding the ink for a predefined time and measuring the extruded weight. Further, to measure the filament width, fluorescent micro-particles (1 µm in diameter) were added to the hydrogel (1.5% XaGMA/ 5% GelMA) and the filaments were printed inside the support bath. This assembly was then directly used to take the micrographs of the filaments using the Zeiss Axio Observer. The filament widths were measured and compared with theoretical filament diameter calculated using the flow continuity equation ^[18]^.

### Aspiration-assisted Bioprinting of Spheroids and its Characterization

A custom-made AAB system was used to bioprint spheroids inside the hydrogels as reported before ^[33,34]^ and characterization of AAB was performed according to the literature ^[22,33]^. To evaluate the bioprinting accuracy and precision of spheroids, first, a support bath was cast into a Petri dish and composite hydrogels (1% XaGMA/ 5% GelMA, 1.5% XaGMA/ 5% GelMA, and 2% XaGMA/ 5% GelMA) were bioprinted inside the support bath in a single layer. After that, hMSCs spheroids were bioprinted inside the XaGMA/GelMA at a predetermined target position on a micrometer calibration ruler. The calibration ruler was placed at the bottom of the Petri dish and the bioprinting steps were recorded by a microscopic camera (Plugable USB Digital Microscope, Plugable Technologies, Redmond, WA, USA) to monitor the target position. For each composite hydrogel group, a total of 10 spheroids were bioprinted and imaged. The images were analyzed using ImageJ (National Institutes of Health, MD, USA) and the accuracy of the printed spheroids was calculated using the following equation:

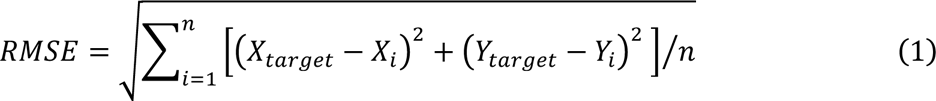

Where *RMSE* was the root mean square error, *n* was the sample size, *X_target_* and *Y_target_* were the coordinates of the spheroid’s target position, *X_i_* and *Y_i_* were the actual coordinates of *i*^th^ spheroid’s position after bioprinting, where *i*=1, 2, 3…*n*. Precision was represented as the square root of the standard deviation.

### IEB with a Silicone Composite

To fabricate a pancreas model, a photo-crosslinkable silicone composite was used. The silicone composite was prepared as per previous research ^[35]^. Briefly, 37.5 g vinyl terminated silicone copolymer and 8 g of mercaptopropyl functionalized silicone copolymer were added to a 100 mL container and mixed in a speed mixer (DAC-330-110 SE, Flacktek Manufacturing, Landrum, SC, USA) at 3,500 rpm for 1 min. Then, 2.5 g of HMDS-treated silica was added to the container and mixed at 3,500 rpm for 5 min. After mixing, 2 g of thixotropic agent was added to the container and mixed at 3,500 rpm for 2 min. Finally, 100 µL of 10% (w/v) 2,2-Dimethoxy-2-phenylacetophenone dissolved in ethanol was added to the container and mixed at 3,500 rpm for 5 min. The mixture was loaded into 3 mL syringe barrels for printing purposes and centrifuged at 4,000 rpm for 4 min to remove bubbles. To accomplish IEB of the pancreas model, head phantom and Matroshka doll, the silicone composite was printed inside a 1-3% (w/v) XaG support bath. Thereafter, IEB of the sacrificial ink made of XaG and different color dyes (Winton Oil Colour, UK) was carried out inside the printed silicone model to replicate the vasculatures and the bile duct of the pancreas.

### IEB of a Breast Tumor Model

First, 5 × 10^5^ /mL ADSC-derived adipose cells were loaded into 1.5% (w/v) XaGMA/ 5% (w/v) GelMA solution and then bioprinted into 1.5% (w/v) XaGMA/ 5% (w/v) GelMA. Next, a sacrificial ink made of 3% (w/v) XaG was printed into XaGMA/GelMA inside the breast tumor model to be used later as a perfusable channel. Then, using AAB, three tumor spheroids were bioprinted inside the breast tumor model around the sacrificial ink within a distance of 200-400 µm. After that, the support bath as well as the breast model was photo-crosslinked, and the sacrificial ink was removed to obtain the perfusable channel. The model was incubated in the media under static conditions for two days before starting the DOX (Selleck Chemicals, TX, USA) drug treatment. DOX (10 and 50 µM) was perfused through the channel at a rate of 0.70 µL/min using ISMATEC peristaltic pumps (Cole Palmer, IL, USA) for 1 or 3 days, and the cancer spheroids were imaged using a ZEISS LSM 880 confocal microscope (Thornwood, NY, USA) and the tumor volume was quantified using ImageJ. For analysis, the automatic threshold method was used to outline each spheroid, allowing for the measurement of the thresholded object consisting of multiple points across all consecutive slices. The volume was calculated by multiplying the slice thickness with the sum of the surface area of all the spheroid outlines.

### Statistical Analysis

All data were presented as mean ± standard deviation. Data were analyzed by GraphPad Prism (Dotmatics, USA) and statistical differences were determined using one-way analysis of variance (ANOVA) with Tukey’s post hoc test, and the analysis, if fulfilling the null hypothesis at *p* ≤ 0.05 (^∗^), *p* ≤ 0.01 (^∗∗^), *p* ≤ 0.001 (^∗∗∗^) and *p* ≤ 0.0001 (^∗∗∗∗^) were considered statistically significant.

## Acknowledgement

This work has been supported by National Science Foundation Award 1914885, National Institute of Dental and Craniofacial Research Award R01DE028614, National Institute of Allergy and Infectious Diseases Award U19AI142733, 2236 CoCirculation2 of TUBITAK award 121C359. We thank Dr. Danny Welch, from University of Kansas, Kansas City, USA for gifting GFP^+^ MDA-MB-231 metastatic breast cancer cells used in the study. The opinions, interpretations, conclusions, and recommendations are those of the author and are not necessarily endorsed by National Science Foundation, National Institute of Dental and Craniofacial Research, National Institute of Allergy and Infectious Diseases Award and TUBITAK. YOY acknowledges the support from The Scientific and Technological Research Council of Turkey (TUBITAK) under the BIDEB/2214-A Doctoral Scholarship Program.

