## Supporting Information for "Nested biofabrication: Matryoshka-inspired Intra-embedded Bioprinting"

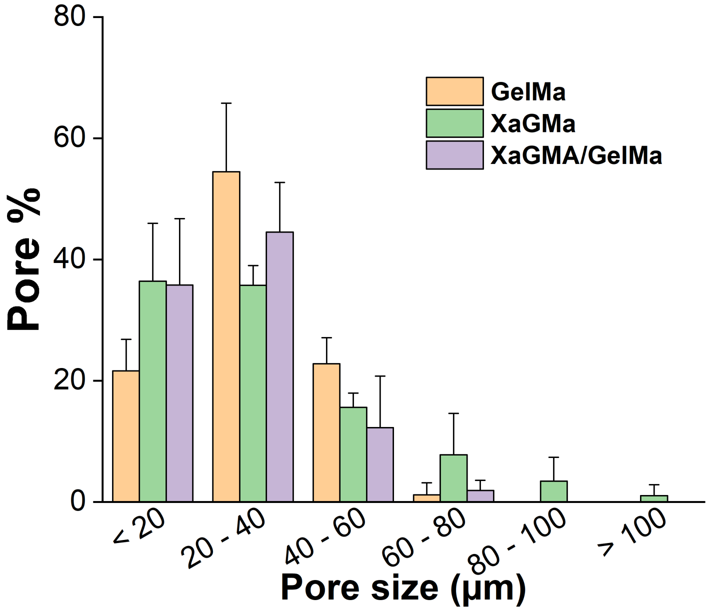


**Figure S1.** The pore size distribution in XaGMA, GelMA and the XaGMA/GelMA composite (*n*=3).

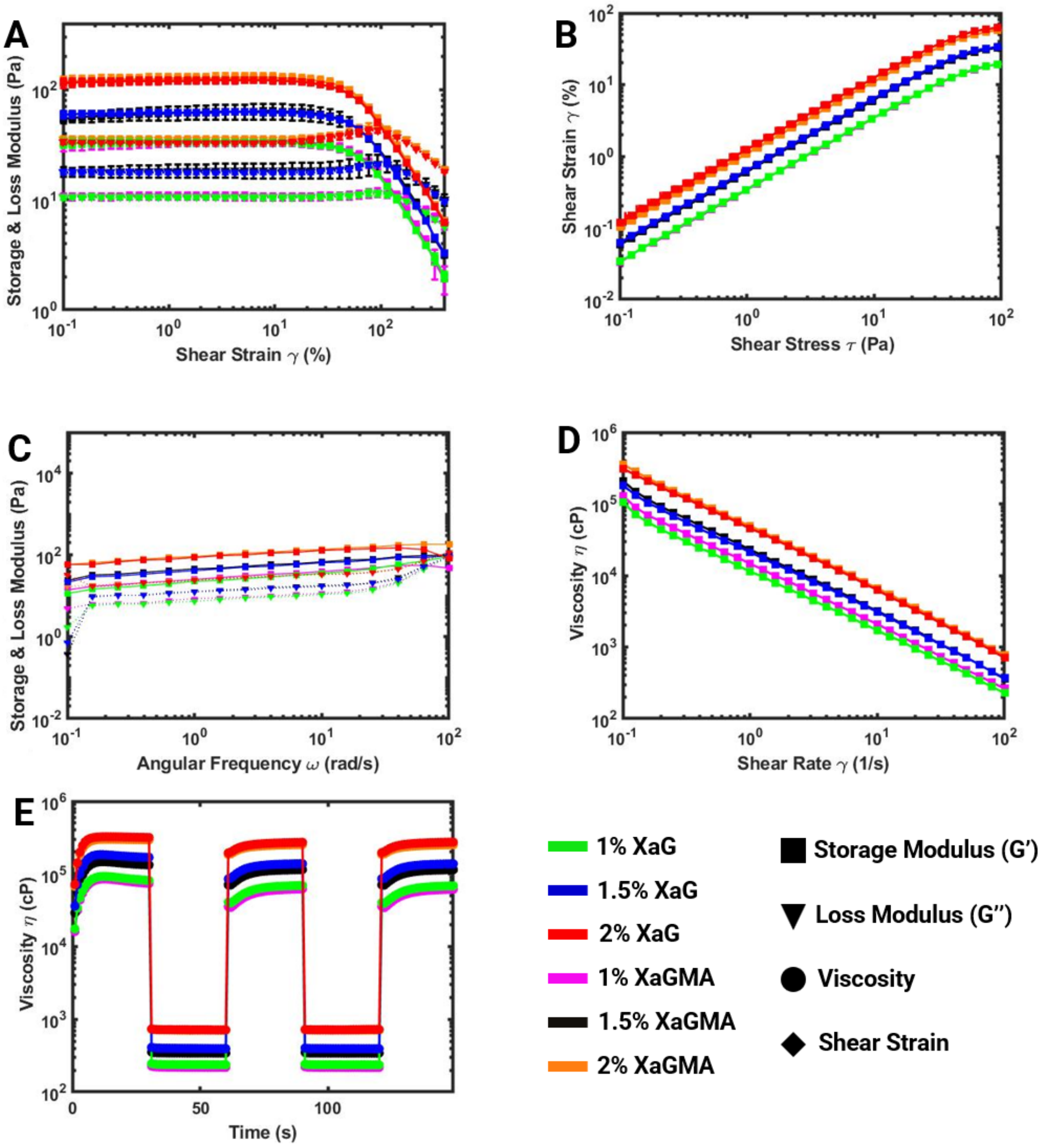


**Figure S2. Rheological analysis of XaGMA.** Rheology of XaG compared with the XaGMA composite, (A) amplitude sweep test, (B) stress vs. strain curve, (C) frequency sweep test, (D) flow curve, and (E) thixotropy test (*n* = 3).

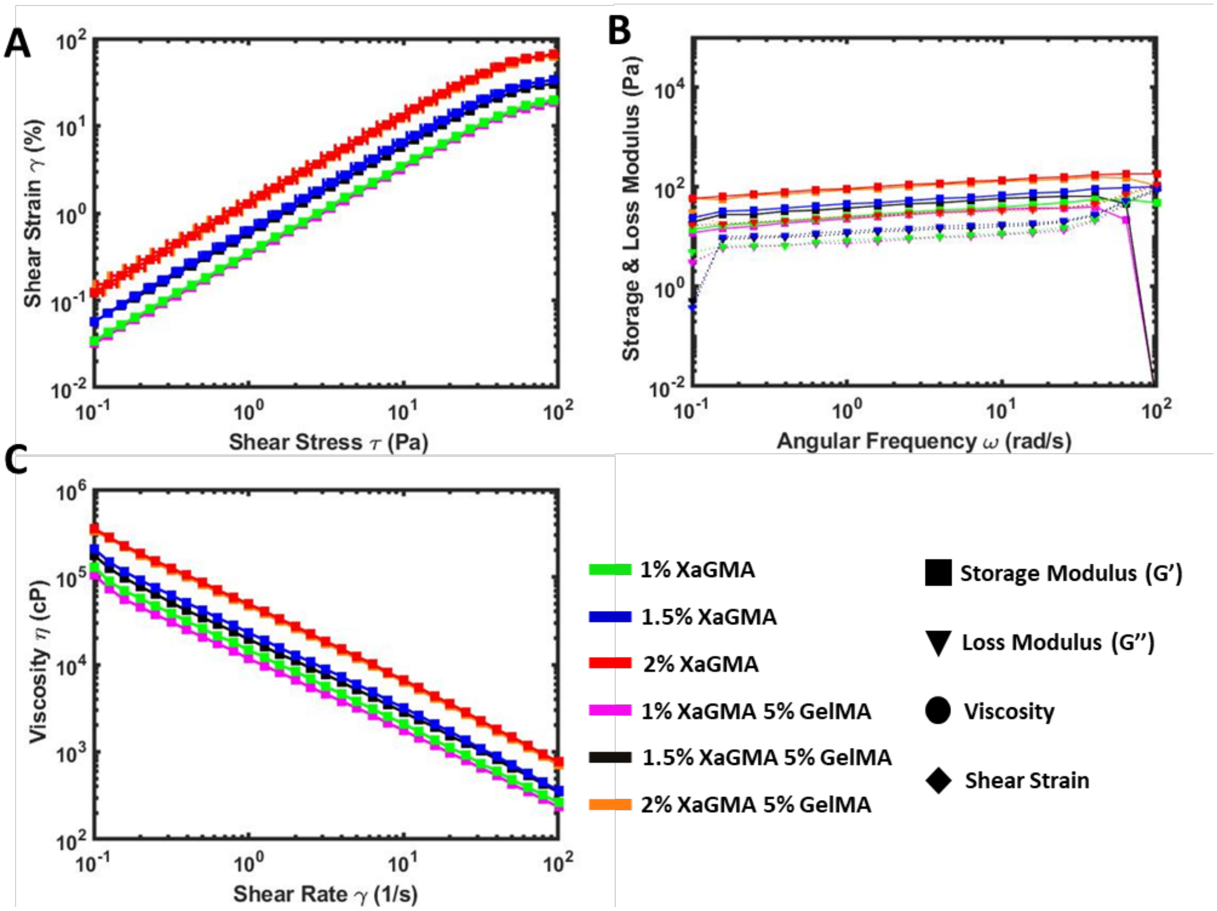


**Figure S3. Rheological analysis of XaGMA/GelMA.** (A) Stress-strain curve, (B) frequency sweep test to validate the elastic nature of the material at low shear strain and an angular frequency ranging from 0.1 to 100 rad $s^{-1}$, and (C) flow sweep test to measure the viscosity of the material under increasing shear rate (*n*=3).

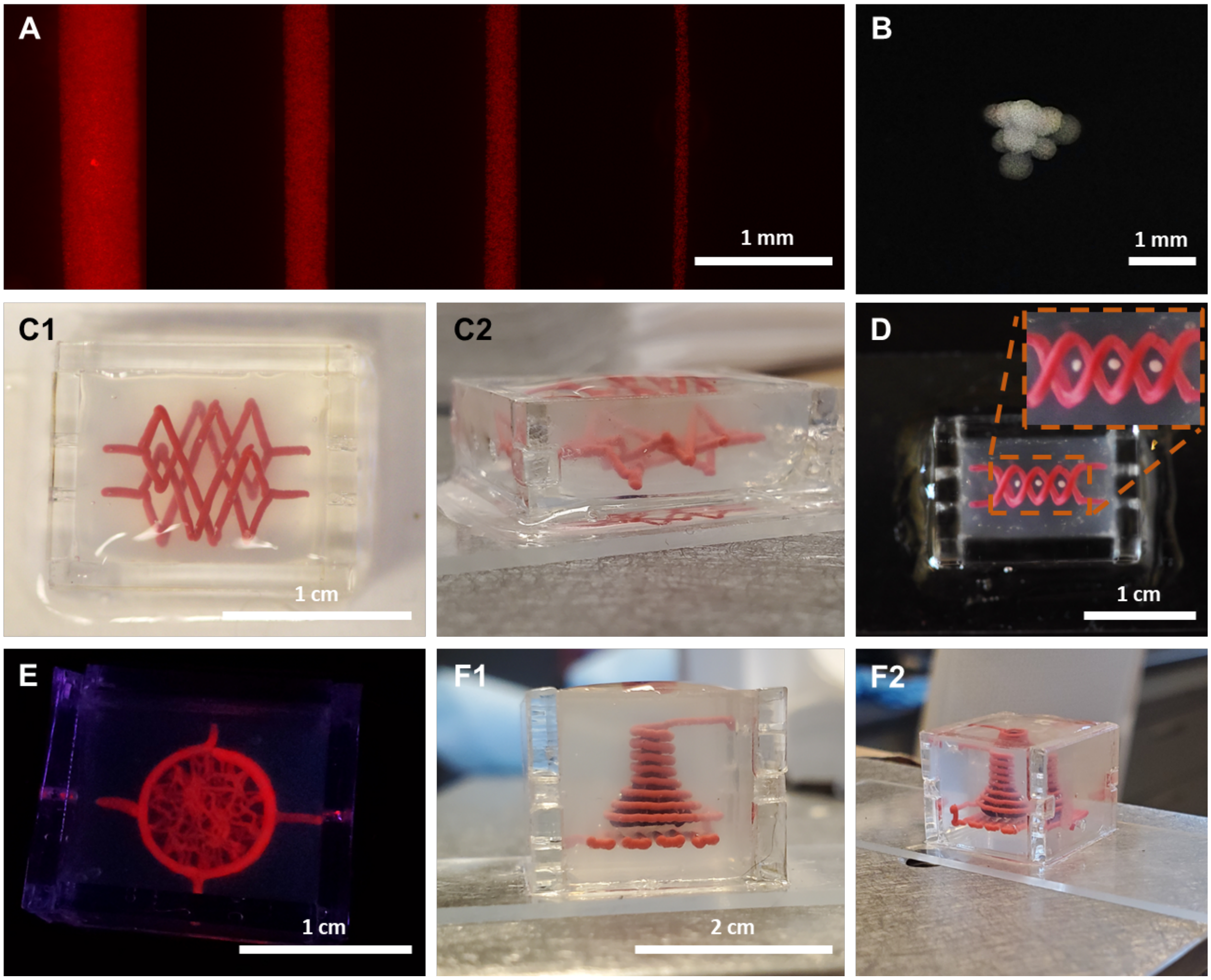


**Figure S4. Embedded printing in XaG.** Embedded printing of (A) filaments ranging from 100 to 600 µm, and (B) multiple spheroids in the shape of a pyramid. Embedded printing of (C1-C2) an intertwined channels design, (D) double helix channels with spheroids and (E) a bi-layer vascular bed design with a circumfential larger channel and smaller vascular channels within it. (F1-F2) Embedded printing of a lung model, where the alvelolar chamber was enclosed by a vasculature.

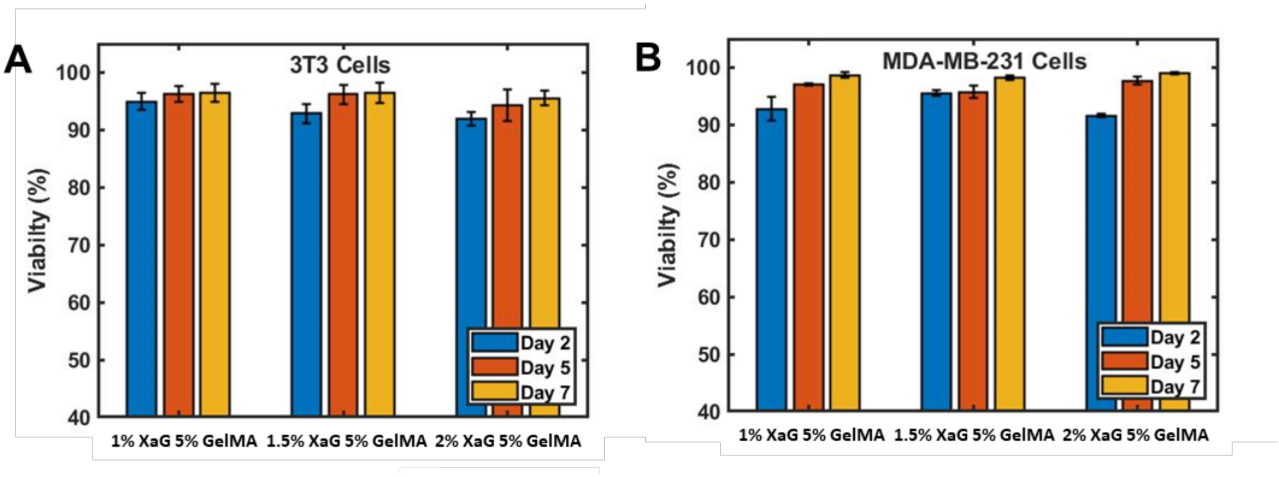


**Figure S5. Biocompatibilty of XaG/GelMA.** (A) Viability of 3T3 and (B) MDA-MB-231 cells on XaG/GelMA composites with different XaG concentrations (*n*=3).

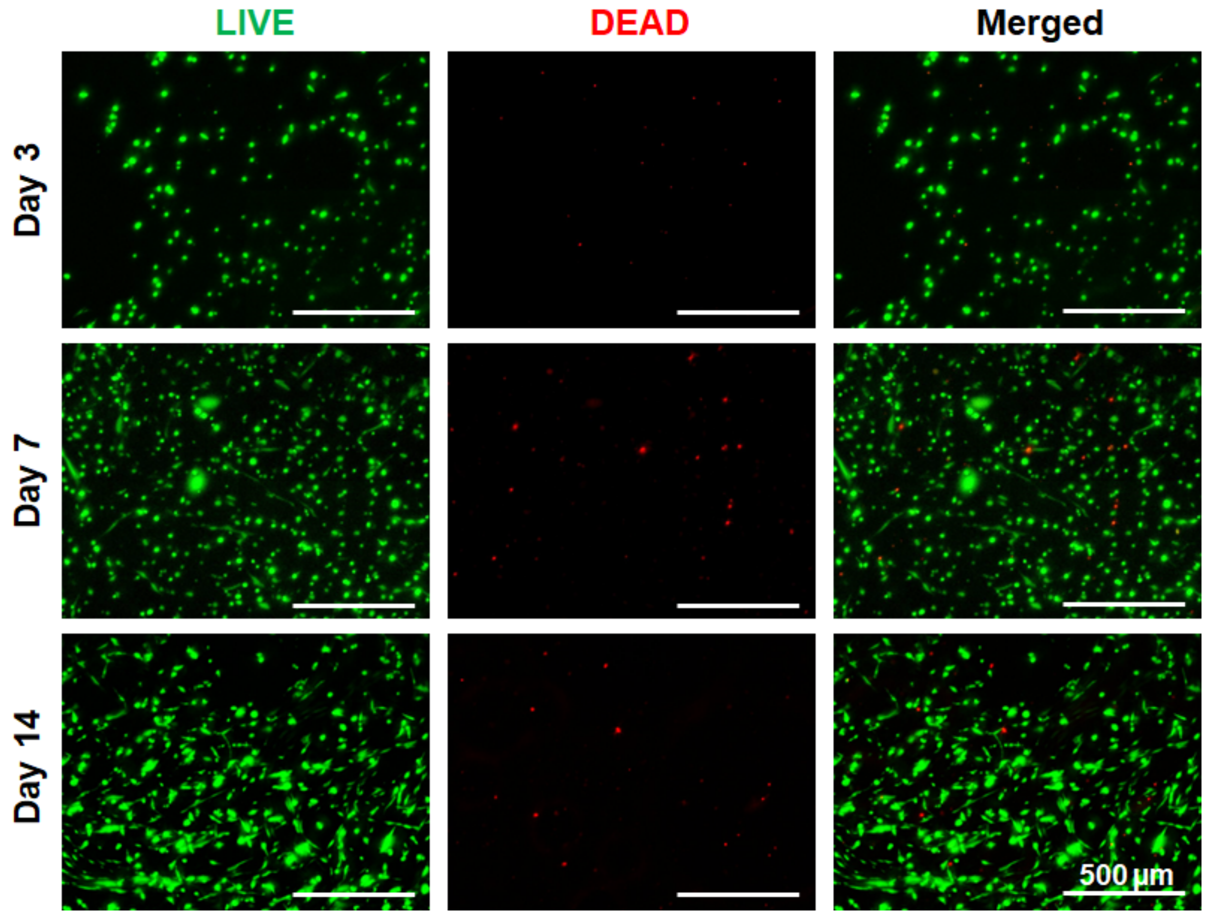


**Figure S6. Viability of ADSCs in XaGMA/GelMA.** LIVE/DEAD images of ADSCs on cast samples at Days 3, 7, and 14.

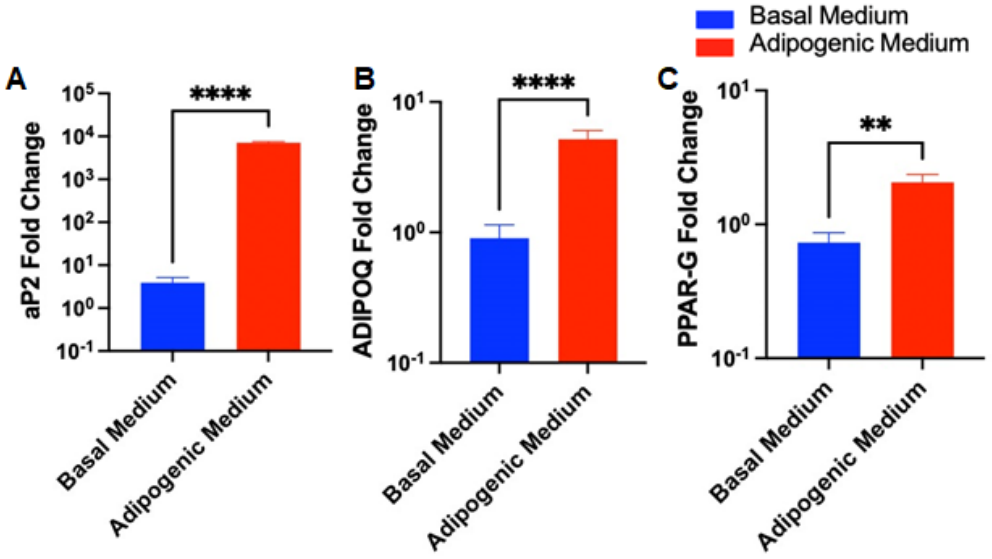


**Figure S7. Gene expression of adipogenic markers using qRT-PCR on Day 15.** (A) Adipocyte fatty acid-binding protein (aP2), (B) adiponectin (ADIPOQ), and Peroxisome proliferator-activated receptor gamma (PPAR-G). Data were presented as mean ± S.D. (*n* = 3; **p ≤ 0.01, ****p ≤ 0.001).

**Table S1.** Primers of the genes used in qRT-PCR study.

| Gene | Forward primer | Reverse primer |
|---|---|---|
| **aP2** | 5'-ATG GGA TGG AAA ATC AAC CA-3' | 5'-GTG GAA GTG ACG CCT TTC AT-3' |
| **PPAR-G** | 5'-TCA GGT TTG GGC GGA TGC-3' | 5'-TCA GCG GGA AGG ACT TTA TGT ATG-3' |
| **ADIPOQ** | 5′-TGA CGA CAC CAA AAG GGC-3′ | 5′-GTG TGT CGA CTG TTC CAT GA-3′ |
| **B-ACTIN** | 5′-GCC CAC ATC TCC ACC TAT GAT-3′ | 5′-GCA GTT CTC GTT GTC CGT CA-3′ |

**Movie Captions:**

**Movie 1:** IEB of a pancreas model (10X playback speed)

**Movie 2:** IEB of a head phantom model (50X playback speed)

**Movie 3:** IEB of a Matryoshka doll (40X playback speed)

**Movie 4:** IEB of a cancer-on-a-chip model (10X playback speed)
